# Toxic effect of 2,2’-bis(bicyclo[2.2.1] heptane) on bacterial cells

**DOI:** 10.1101/2020.01.21.913699

**Authors:** A. G. Kessenikh, L. S. Yaguzhinsky, M. V. Bermeshev, V. G. Pevgov, V. O. Samoilov, S. V. Shorunov, A. L. Maksimov, L. S. Yaguzhinsky, I. V. Manukhov

## Abstract

Here we present the study of the genotoxic effect of a 2,2’-bis(bicyclo[2.2.1]heptane) (BBH), which is promising as a fuel component for liquid rocket engines. The use of *Escherichia coli lux*-biosensors showed that in addition to DNA damage causing SOS-response, there is also an oxidative effect on cells. The greatest toxicity is determined by the mechanism of formation of superoxide anion radical and is detected by the lux biosensor *E. coli* pSoxS-lux, in which the genes of bacterial luciferases are transcriptionally crosslinked with the promoter of the *soxS* gene. It is assumed that the oxidation of BBH leads to the formation of reactive oxygen species, which should give the main contribution to the toxicity of this substance.

## Introduction

Norbornane and its non-saturated derivatives are commonly used in the production of rubber, epoxides, medicinal compounds and perfumes [1, 2]. Notably, thermotechnical characteristics of other strained hydrocarbons made them attractive for high-performance combustion applications. Strained 2,2’-bis(bicyclo[2.2.1]heptane) (BBH) compound is a promising as a fuel component for liquid rocket engines. It is assumed that BBH is comparable with the unsymmetrical dimethylhydrazine (UDMH) in terms of specific impulse efficiency, but can be significantly less toxic to the environment and personnel working with rocketry. For the purpose of this study, BBH which consists of two strained structures made of 14 carbons and 22 hydrogens (Fig. 1), was synthesized from 5-vynil-2-norborene [3-5]. Here we present the data describing toxicity of BBH, which we had obtained by utilizing bacterial *lux*-biosensors.

**Figure 1.**
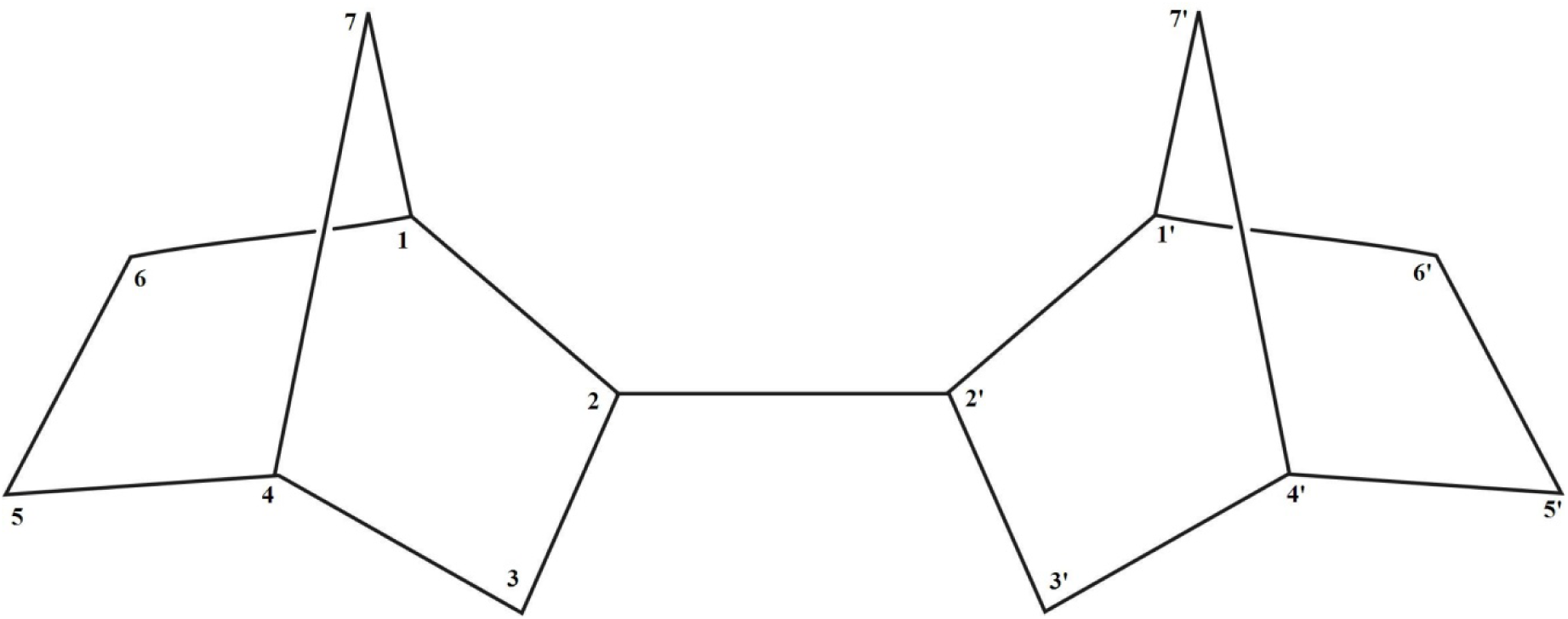
Structure of 2,2’-bis(bicyclo[2.2.1]heptane) molecule [3, 4].

*Lux*-biosensors are living *Escherichia coli* cells transformed with hybrid plasmids, containing *luxCDABE* genes of *Photorhabdus luminescense* placed under control of various stress-responsive promoters, responsible for increasing the cells luminescence in occurrence of toxicants in the environment [6-9]. Similarly designed study of UDMH toxicity has been completed earlier [13, 14].

In this study, the genotoxicity of BBH was evaluated using specific lux-biosensors of *E. coli* MG1655 cells with hybrid plasmids pAlkA-lux, pOxyS-lux, pSoxS-lux and pColD-lux, reacting to DNA alkylation, oxidative damage by hydrogen peroxide, superoxide anion radicals and DNA damages that cause an SOS response, respectively. Were compared of threshold concentrations effect of BBH and UDMH on different stress-responsive promoters.

## Materials and Methods

### Bacterial strains and plasmids

The cells of *Escherichia coli* K12 strain MG1655 F^-^ *ilv*G *rfb*-50 *rph*-1 were combined with plasmids pAlkA-lux, pColD-lux, pOxyR-lux and pSoxS-lux [8-12] that were built using a promoterless plasmid backbone pDW201 [7]. *E. coli* MG1655 cells transformed with pXen7 plasmid constitutively expressing *luxCDABE* genes [15] were employed as non-inducible control.

### Media and Culturing conditions

The cells of *E. coli* were grown at 30°C in LB broth supplemented with ampicillin (100 µg/ml), in aerated conditions until early exponential phase.

### Measurement of the intensity of bioluminescence

Cell were prepared by overnight cultivation at 30°C with aeration at 200 rpm in LB media, then diluted 1:100 in LB media, grown till reaching *OD*_*600*_ = 0.1-0.2, which corresponds to early/mid-logarithmic phase. These cells were sampled into the 200-μl subcultures in separate tubes, then 10 μl of tested compound (BBH or control) were added. Cells were grown without shaking at ∼30°C, with repetitive direct measurements of total bioluminescence (in RLU, relative light units) using “Biotox-7” (LLC EKON, Russia) or plate luminometer LM–01A (Immunotech). Visible enhancement of bioluminescence had been detected in 15-20 minutes of incubation, which roughly corresponds to the time necessary for biosynthesis of luciferase. Maxima of bioluminescence were observed after 40-60 minutes of exposure for all plasmids except one with the promoter P_colD._ In case of latter construct, maximum bioluminescence was detected in 90 minutes post exposure. The magnitude of the response was 50-100 times and was dependent on concentration of toxic agent. Each experiment has been performed in biological triplicates, with each triplicate independently measured five times.

### Chemical

All chemicals were of analytical purity. Hydrogen peroxide was obtained from the firm “Ferraine”. Mitomycin C, N, N ‘-dimethyl-4,4’ - dipyridyl dichloride (paraquat), methyl methanesulfonate obtained from Sigma Chemical Co. All test solutions were prepared immediately before use. The investigated compound 2,2 ‘ - bis (bicyclo[2.2.1]heptane was synthesized by the Diels-alder reaction from 5-vinyl-2-norbornene and Dicyclopentadiene according to [16] with the subsequent stage of exhaustive hydrogenation of the cycloadduct in methanol on a Pd/C catalyst (1%) with hydrogen (25°C, 20 ATM, 24 h) [1, 5].

## Results

At the first stage, the ability of BBH to alkylate DNA was investigated. To do this, *E. coli* MG1655 cells transformed by pAlkA-lux plasmid were grown to DO=0.1, then they were added to different concentrations of BBH. The mixtures were incubated at room temperature without aeration for 3 hours with periodic luminescence measurement. Figure 2 shows the luminescence kinetics of *E. coli* MG1655 (pAlkA-lux), after the addition of BBH. The pAlkA-lux plasmid contains *lux* genes under the control of the Alka gene promoter, therefore when alkylating agents occur in the sample the luminescent response increases. Dilutions of the alkylating substance methyl methanesulfonate (MMS) were used as a positive control.

**Figure 2.**
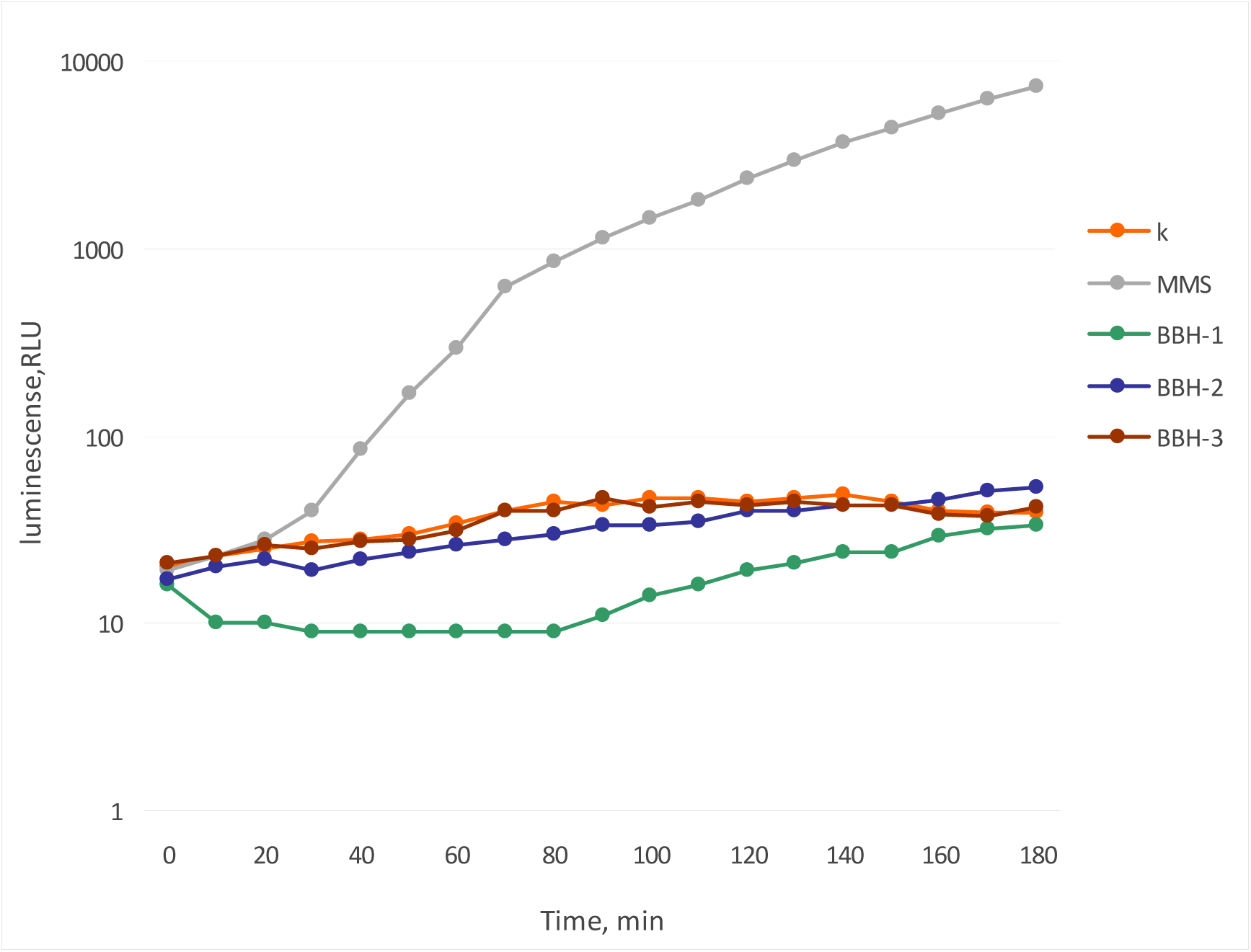
Luminescence of *E. coli* MG1655 pAlkA-lux cells after BBH addition depending on incubation time. k-control cells without toxicant addition mms - added MMS to final concentration of 100 μM BBH-1-added BBH to final concentration of 100 g/l, BBH-2 – 10 g/l, BBH-3 – 1 g/l

As can be seen from the data shown in figure 2, none of the tested concentrations of BBH does not cause an alkylating effect (luminescence of the Alka-lux biosensor does not increase). At a maximum concentration of 100 g/l (10%) BBH has a cytotoxic effect, causing a slight decrease in the background luminescence of cells (about 2-3 times).

Figure 3 shows the measurement of the luminescence kinetics of *E. coli* MG1655 containing pOxyR-lux plasmid, after the addition of BBH. Hydrogen peroxide was used as a positive control. The cells were incubated at room temperature for 2 hours with periodic luminescence measurement.

**Figure 3.**
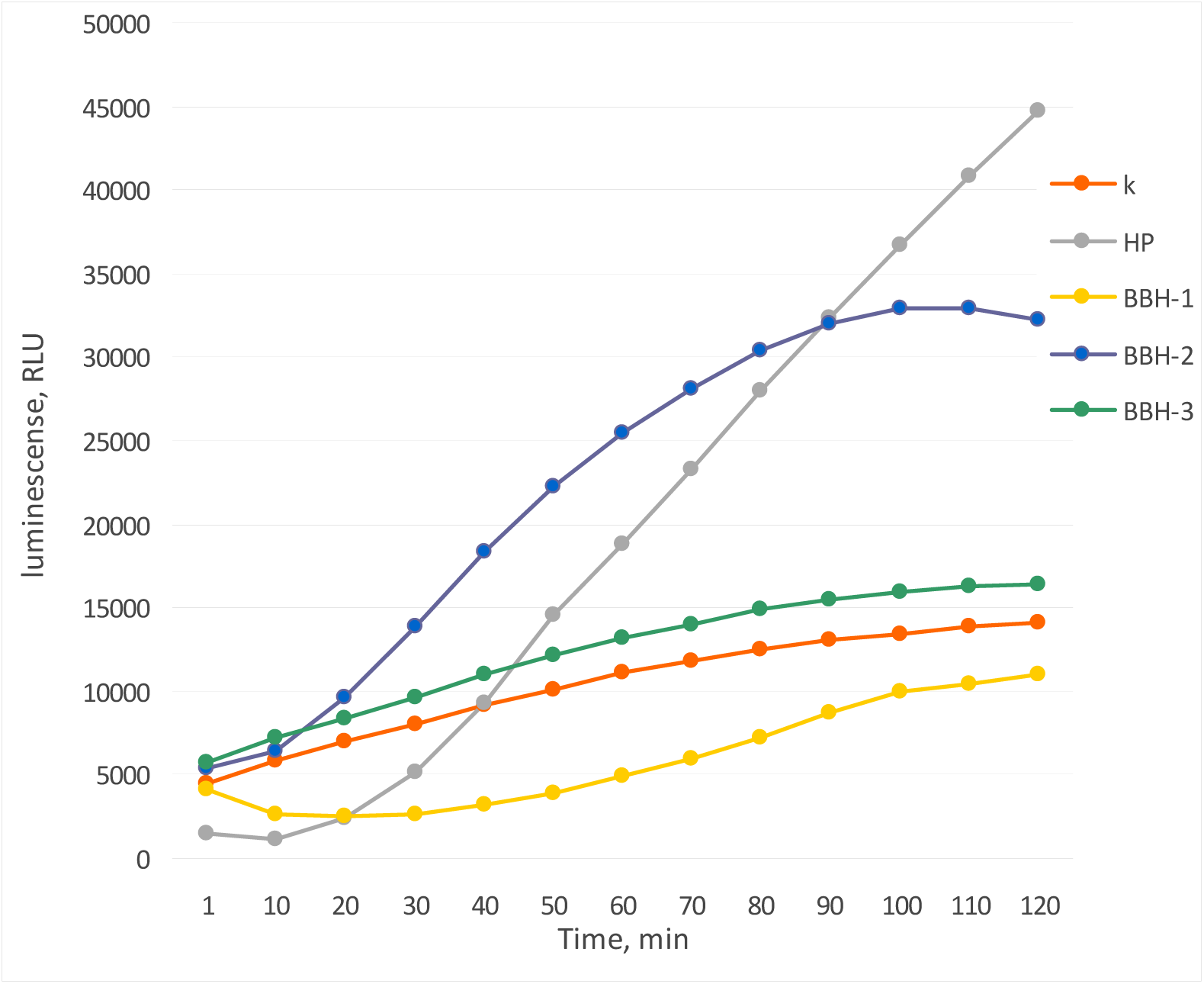
Luminescence of *E. coli* MG1655 pOxyR-lux cells after BBH addition depending on incubation time. k - control cells of *E. coli* MG1655 pOxyR-lux without toxicant addition HP - added hydrogen peroxide to a final concentration of 1 mM. BBH-1-added BBH to final concentration of 100 g / l, BBH -2-10 g / l, BBH -3 – 1 g/l

As can be seen from the data presented in the figure 3 there is oxidative stress caused by hydrogen peroxide. This type of cell damage can lead to modifications of DNA nitrogenous bases, leading to an increase in the rate of mutagenesis. The maximum effect is achieved at a concentration of BBH corresponding to 1% content in water (fig. 3, curve BBH-2), a lower concentration does not cause a significant increase in luminescence, and a higher has a General toxic effect on cells, leading to a decrease in the base level of luminescence.

Figure 4 shows the measurement of luminescence kinetics of *E. coli* MG1655 with pCold-lux plasmid, after the addition of BBH. As a positive control, were used the antibiotic mitomycin C, which forms crosslinking with DNA. As a result, there is a stop of the replication fork, the formation of single-stranded DNA sites and, as a consequence, SOS response. The cells were incubated at room temperature without aeration for 5 hours with periodic luminescence measurement.

**Figure 4.**
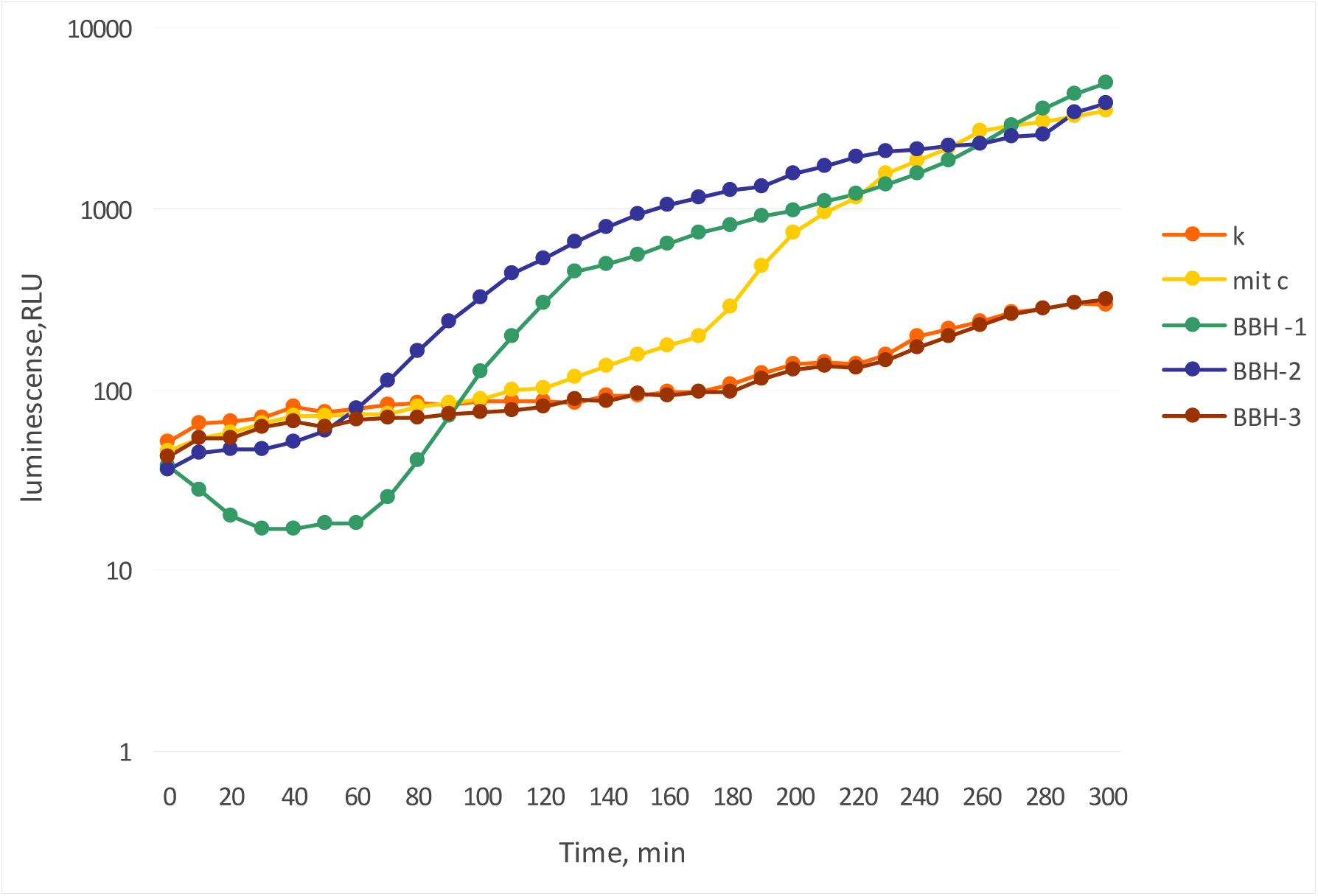
Luminescence of *E. coli* MG1655 pColD-lux cells after BBH addition depending on incubation time. k-*E. coli* MG1655 pColD-lux control cells without toxicant addition mit c - added mitomycin C to the final concentration of 10 μM. BBH-1 - added BBH to final concentration of 100 g / l, BBH-2 - 10 g / l, BBH-3 – 1 g/l

As can be seen from the data shown in the figure, the addition of BBH in concentrations of 10% and 1% leads to a high level of DNA damage, which causes an SOS response. The maximum possible response amplitude of an E. coli biosensor (pColD-lux) is about three orders of magnitude [8]. In this experiment, we see the maximum activation of the biosensor about 20 times when incubated with BBH at a concentration of 10g/l.

Then it was investigated the appearance of a superoxide anion radical in cells during incubation in the presence of BBH. For these purposes, *lux* genes under the control of the Psox promoter were used. Figure 5 shows the luminescence kinetics of *E. coli* MG1655 pSoxS-lux cells incubated with BBH at room temperature without aeration for 4 hours. As a standard inductor for a Psox promoter is usually used paraquat – a substance that, when ingested, leads to occurrence of superoxide anion radical as a result of reactions with quinones of the respiratory chain. Hydrogen peroxide does not directly react with the SoxR protein, but causes lipid peroxidation and oxidation of a number of proteins, which in turn disrupts the respiratory chain, which eventually leads to increase in the superoxide anion radical pool in the cell [8, 17]. Thus, H_2_O_2_ can also be used as a positive control to *E. coli* MG1655 pSoxS-lux..

**Figure 5.**
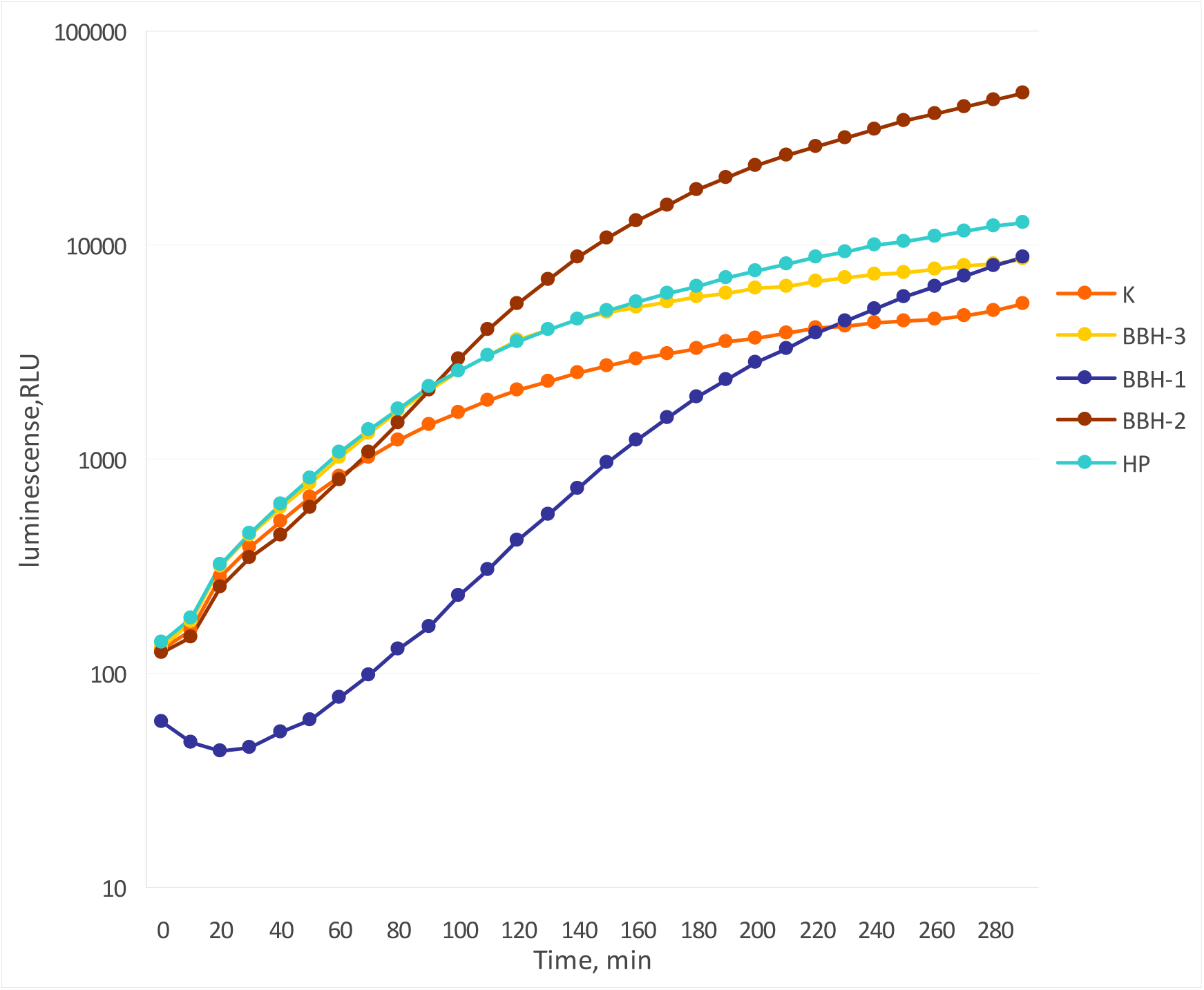
Luminescence of *E. coli* MG1655 pSoxS-lux cells after BBH addition depending on incubation time. k - control cells of *E. coli* MG1655 pSoxS-lux without toxicant addition HP - added hydrogen peroxide to a final concentration of 1 mM. BBH-1 - added BBH to final concentration of 100 g / l, BBH-2 - 10 g/l, BBH-3 – 1 g/l

As can be seen from figure 5, activation of the PsoxS promoter occurs when BBH is added at all concentrations from 1 to 100 g/l. On the basis of the data obtained in the experiments we can suggest that BBH causes the appearance of a superoxide anion radical in the cell.

Table 1 shows the threshold values of activation of stress promoters when UDMH and BBH appear in the medium. As “control” data are given for standard, promoter-specific toxicants inducing a noticeable (1.5-2 times) effect of bioluminescence enhancement of lux-biosensors. Mitomycin C induces damage in DNA, hydrogen peroxide causes the oxidative stress, paraquat used as a generator of superoxide ion radicals witch are induce oxidative stress, methylnitronitrosoguanidine (N-methyl-N’-nitro-N-nitrosoguanidine H2NC(=NH)NHNO) (MNNG) alkylates DNA.

**Table 1.**
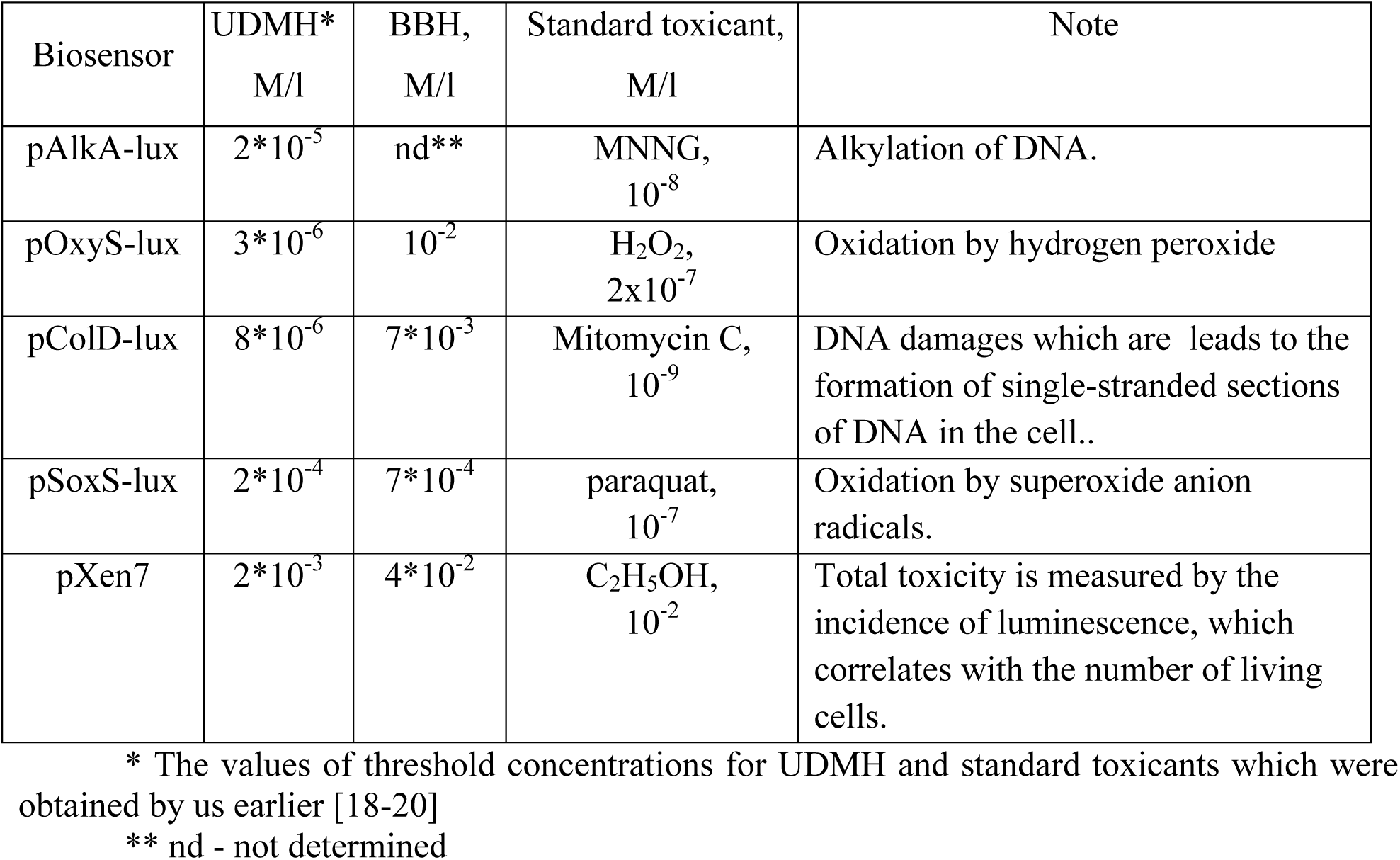
Threshold concentrations for lux biosensors.

| Biosensor | UDMH*<br>M/l | BBH,<br>M/l | Standard toxicant,<br>M/l | Note |
| --- | --- | --- | --- | --- |
| pAlkA-lux | $2 \cdot 10^{-5}$ | nd** | MNNG,<br>$10^{-8}$ | Alkylation of DNA. |
| pOxyS-lux | $3 \cdot 10^{-6}$ | $10^{-2}$ | $\text{H}_2\text{O}_2$,<br>$2 \cdot 10^{-7}$ | Oxidation by hydrogen peroxide |
| pColD-lux | $8 \cdot 10^{-6}$ | $7 \cdot 10^{-3}$ | Mitomycin C,<br>$10^{-9}$ | DNA damages which are leads to the formation of single-stranded sections of DNA in the cell.. |
| pSoxS-lux | $2 \cdot 10^{-4}$ | $7 \cdot 10^{-4}$ | paraquat,<br>$10^{-7}$ | Oxidation by superoxide anion radicals. |
| pXen7 | $2 \cdot 10^{-3}$ | $4 \cdot 10^{-2}$ | $\text{C}_2\text{H}_5\text{OH}$,<br>$10^{-2}$ | Total toxicity is measured by the incidence of luminescence, which correlates with the number of living cells. |

\* The values of threshold concentrations for UDMH and standard toxicants which were obtained by us earlier [18-20]
\*\* nd - not determined

## Discussion

It is known that as a result of oxidation of a number of hydrocarbon compounds by bacterial cells, oxidative stress occurs [21, 22].

According to the literature, BBH must be oxidized by oxygen, as well as all hydrocarbons by a free-radical chain mechanism [23]. In the same work it is shown that under intense light one of the products of strained compounds destruction is hydroxyl radical, this is shown by the example of cyclopropane. The BBH molecule includes two elements with strained bonds, which determine a higher energy release in the oxidation process compared to conventional non-strained hydrocarbons. Comparison of ΔH≠ (in Kcal/mol) obtained for ring opening of cycloalkanes [24] shows that the ring strain energy is 3.57 times higher in cycloalkanes compared to cyclopentane. The radical mechanism of oxidation along with the increased reaction energy suggests that the toxic effect of this type of compounds should be determined mainly by reactions of formation of reactive oxygen species. In this regard, it should be noted that according to our experiment, the toxicity of BBH, determined by the appearance of a superoxide anion radical in cells, is close to that of UDMH, despite the fact that the tests were conducted with an undispersed form of the product insoluble in water, whose contacts with biological objects are sharply reduced and are determined only by the interface of the phases. Under natural conditions, deep dispersion of products is completely absent during fuel spillage, including in a humid environment. Thus, the chosen experimental conditions achieve convergence with the conditions of accidental fuel spill.

The genotoxic effect of BBH definitely takes place and is expressed by cell damages that activates the following defense systems: SOS response, oxyRS regulon, and soxRS regulon. Activation of SOS response occurs only at very high concentrations of BBH in the medium (threshold concentration is about 1 g / l). The test using *E. coli* MG1655 pColD-lux is more sensitive to genotoxic agents than SOS chromotest [25], which is used in toxicology along with the Ames test [26, 27] to determine the rate of mutagenesis. These tests correlate well with each other [28], but may underestimate the effect of alkylating compounds on the rate of mutagenesis [29]. In the present work, using the *E. coli* MG1655 pAlkA-lux biosensor, it was shown that during incubation of cells with BBH, DNA alkylation is not observed and, obviously, alkylating compounds do not appear in the medium. We can proposed that the oxidation of BBH in the medium in the presence of *E. coli* cells leads to the appearance of alkyl radicals or alkyl hydroperoxides, which can lead to DNA damages. Thus, the mechanism of genotoxic action of BBH is fundamentally different from the action of UDMH, which is determined by alkylating derivatives, primarily nitrosodimethylamine [14] and superoxide-anion radicals arising from the oxidation of UDMH with atmospheric oxygen [13].

## ACKNOWLEDGMENTS

The work was financially supported by the Ministry of Higher Education and Science of Russia (Project Unique Identifier RFMEFI60417×0181, Agreement No. 14.604.21.0181 of 26.09.2017)

